# Optimal Protein Allocation Controls the Inhibition of GltA and AcnB in *Neisseria gonorrhoeae*

**DOI:** 10.1101/2024.05.22.595331

**Authors:** Nabia Shahreen, Niaz Bahar Chowdhury, Rajib Saha

**Affiliations:** Department of Chemical and Biomolecular Engineering, University of Nebraska-Lincoln

**Keywords:** Optimal protein allocation, constraint-based modeling, energy production, acetate overflow, *Neisseria gonorrhoeae*

## Abstract

*Neisseria gonorrhea* (Ngo) is a major concern for global public health due to its severe implications for reproductive health. Understanding its metabolic phenotype is crucial for comprehending its pathogenicity. Despite Ngo’s ability to encode TCA cycle proteins, GltA and AcnB, their activities are notably restricted. To investigate this phenomenon, we used the iNgo_557 metabolic model and incorporated a constraint on total cellular protein content. Our results indicate that low cellular protein content severely limits GltA and AcnB activity, leading to a shift towards acetate overflow for ATP production, which is more efficient in terms of protein usage. Surprisingly, increasing cellular protein content alleviates this restriction on GltA and AcnB and delays the onset of acetate overflow, highlighting protein allocation as a critical determinant in understanding Ngo’s metabolic phenotype. These findings underscore the significance of Ngo’s metabolic adaptation in light of optimal protein allocation, providing a blueprint to understand Ngo’s metabolic landscape.

## Introduction

*Neisseria gonorrhoeae* (Ngo), the causative agent of gonorrhea, is one of the most prevalent sexually transmitted infections globally, affecting approximately 87 million people each year, including 1.6 million within the United States (Kreisel *et al*. 2021). This pathogen’s persistence and increasing resistance to antibiotics pose a substantial public health challenge (Chinemerem Nwobodo *et al*. 2022). Ngo’s ability to persist in the human host and to evade the immune system emphasizes the need to understand its unique metabolic phenotype.

Despite having the ability to encode a complete tricarboxylic acid (TCA) cycle, previous studies revealed Ngo’s puzzling phenotype of restricting crucial TCA cycle enzymes, such as GltA and AcnB (Potter and Criss 2023a).Additionally, it does not have the ability to encode phosphofructokinase (PFK). To probe deeper into metabolic anomaly in the TCA cycle, the application of systems biology, particularly, genome-scale metabolic models (GEMs), provides a promising avenue, since it has been proven to answer unusual bacterial phenomena (Chowdhury *et al*. 2023). Previous works has shown that integrating protein allocation constraints within GEMs can significantly enhance our understanding of bacterial metabolism, reflecting more accurately the *in vivo* conditions that might influence metabolic functionality (O’Brien *et al*. 2013; Chen and Nielsen 2019; Chowdhury, Alsiyabi and Saha 2022).

Thus, in this study, we applied a total protein capacity constraint to the recently published iNgo_557 GEM to test the hypothesis that limitations in protein availability affect the activity of TCA cycle enzymes - GltA and AcnB. Our findings indicate that limited protein resources are crucial in suppressing the function of GltA and AcnB enzymes. Notably, by adjusting protein levels within the model, we observed that increased protein availability could relax these restrictions, allowing higher flux through GltA and AcnB and reducing the organism’s reliance on less efficient metabolic pathways such as acetate overflow. These results not only elucidate the significant impact of protein allocation on metabolic pathways but also provide valuable insights into potential metabolic vulnerabilities of Ngo, thus enhancing our understanding on how Ngo manages to sustain itself under the nutrient-limited and immune-rich environment of the human host.

## Materials and Methods

### Initial Model Setup, Data Integration, and Reaction Bounds Modification

The study utilized the previously published Ngo genome-scale metabolic model, iNgo_557 (Potter *et al*. 2023). Initial preparations involved adding pathway information and making the model irreversible (https://github.com/ssbio/Gc_protein_constrained). Following that filtering the model to identify reactions with gene-protein-reaction (GPR) associations. For these reactions, enzyme kinetics (*K*_*cat*_) were sourced predominantly using *DLKcat* (Li *et al*. 2022a), which provides estimated enzyme turnover rates based on homology. Molecular weights (MW) for these enzymes were also calculated using amino acid sequences derived from the KEGG database. Additionally, to accurately reflect the *in vivo* environment, reaction bounds were adjusted according to minimal defined medium (MDM) (Potter *et al*. 2023), ensuring that the simulated nutrient availabilities aligned with the physiological conditions under which *N. gonorrhoeae* thrives.

### Handling Data Gaps with Monte Carlo Simulations and Model Selection

Due to the unavailability of *K*_*cat*_ values for 373 reactions and MW data for 12 reactions, Monte Carlo simulations were conducted to stochastically generate 100 ensemble models (Figure 1A and 1B). These simulations predicted missing *K*_*cat*_ and MW values, filling the gaps necessary for a comprehensive metabolic analysis (Figure 1C and 1D). Each of the 100 simulations provided a unique set of *K*_*cat*_ and MW values for the missing data, which were then integrated into the iNgo_557 model to create 100 ensemble GEMs. For each of these 100 modified GEMs, the cellular protein content 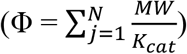 was calculated. Finally, the model with the lowest overall Φ value was selected for further analysis, as it represented the most protein-efficient configuration under protein-limited conditions.

**Figure 1.**
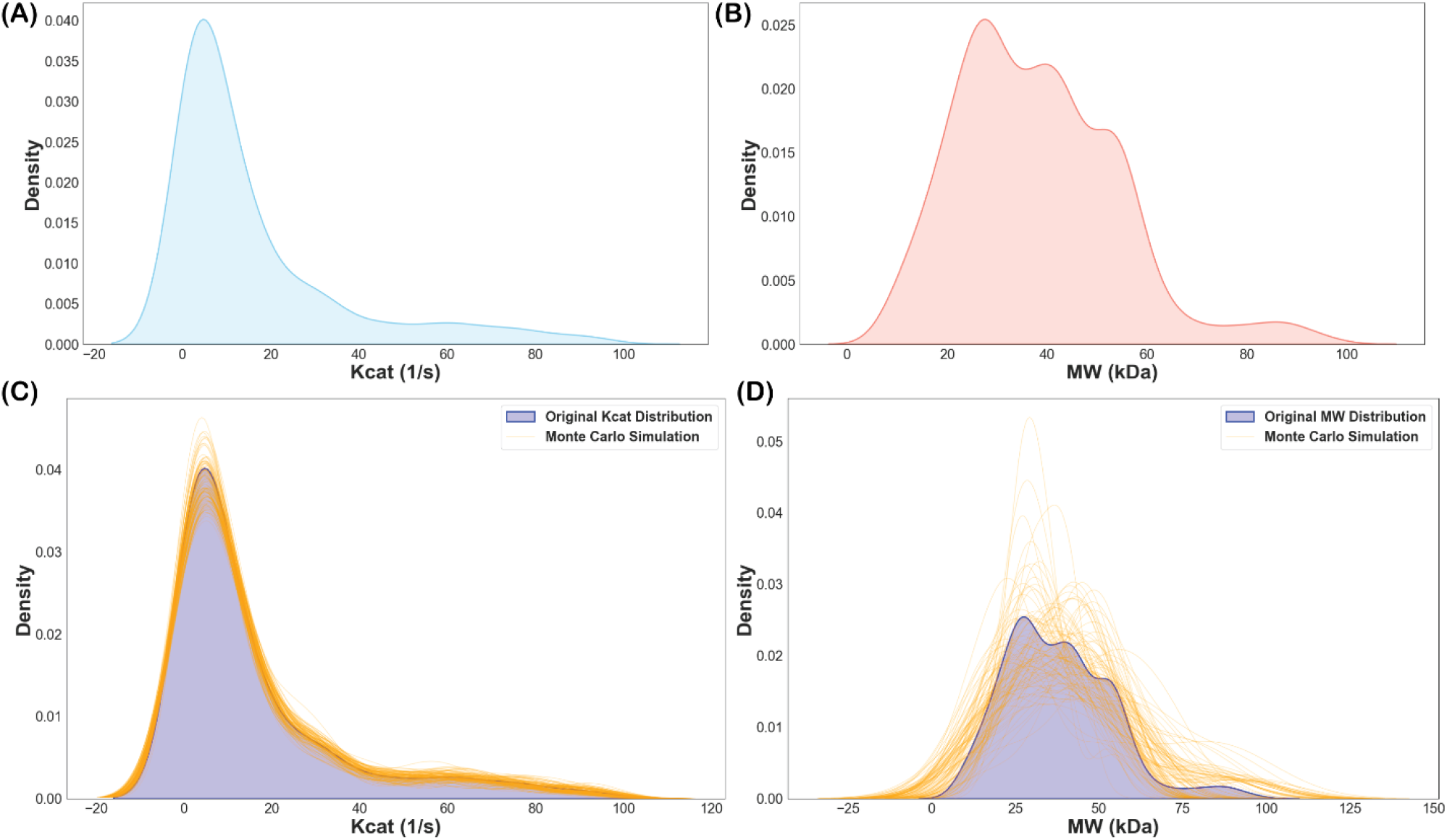
Generation of missing enzyme turn-over rate (*K*_*cat*_) and molecular weight (MW) for missing enzymes. A) Distribution of *K*_*cat*_ for known enzymes. B) Distribution of MW for known enzymes. C) Generated *K*_*cat*_ for 373 missing enzymes for 100 ensembles run using Monte Carlo simulation. D) Generated MW for 12 missing enzymes for 100 ensembles run using Monte Carlo simulation.

### Flux Balance Analysis and Comparative Metabolic Flux Profiling

Next, we employed a sequential optimization strategy within the confines of Flux Balance Analysis (FBA). Initially, the model was tasked with maximizing biomass production. Once the optimal biomass growth rate was established, we fixed this growth rate as a constraint, to maximize acetate production under different glucose uptake while generating fluxes for all the reactions in the model. Moreover, the ATP synthase (ATPS) reaction was not connected to the TCA cycle through the succinate dehydrogenase (SDH) which was resolved by adding an additional constraint.

This configuration was employed for the two distinct protein contents, 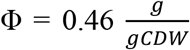 (lower protein content) and 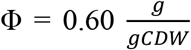 (higher protein content) to assess the flux distribution in different pathways. Following is the mathematical formulation of this optimization problem:

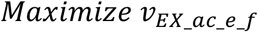

*Subject to*

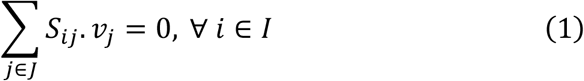

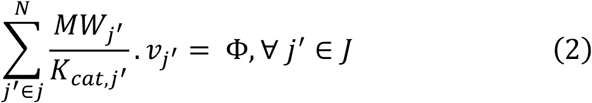

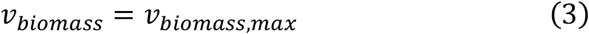

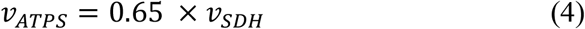

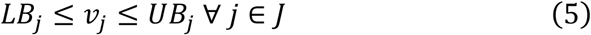

Here, *I* and *J* are the sets of metabolites and reactions in the model, respectively. *S*_*ij*_ is the stoichiometric coefficient of metabolite *i* in reaction *j, j′* is the set of reactions for which GPR is available, Φ is total cellular protein content, and *v*_*j*_ is the flux value of reaction *j*. Parameters *LB*_*j*_ and *UB*_*j*_ denote the minimum and maximum allowable fluxes for reaction *j*, respectively.

Subsequent analysis involved combining the flux distributions for each metabolic pathway. This comparative analysis between the two protein conditions highlighted significant metabolic shifts in revealed critical insights into the metabolic adjustments of Ngo, especially concerning the TCA cycle and acetate overflow pathways.

### Computational Tools

Simulations were conducted using COBRApy, a Python library designed for computational modeling of biological networks, with Gurobi solver for the linear programming tasks. The entire process from data integration to simulation was automated using custom Python scripts, enhancing reproducibility and computational efficiency.

## Results and Discussion

### Limited Protein Availability and Its Impact on TCA Cycle Restriction

In this work, we presented a protein-constrained genome-scale metabolic (pcGEM) model of Ngo to test the hypothesis that optimal proteome allocation influences its unique metabolic phenotype. This includes suppression of the first three steps of the TCA cycle, despite its ability to encode the necessary proteins (Potter and Criss 2023b). To reconstruct the pcGEM, we selected a recently published GEM of Ngo, iNgo_557 (D. *et al*. 2023). We incorporated the pathway information for all the reactions (Dataset S1) and subsequently added unit-flux protein cost data for reactions that has GPR in the model (Chen and Nielsen 2019). The protein cost unit-flux is the ratio of molecular weight (MW) and enzyme turnover rate (*K*_*cat*_) of the enzyme that catalyzes a reaction. Due to the scarcity of experimental *K*_*cat*_ of Ngo, we used *DLKcat* (Li *et al*. 2022b) to predict it (Dataset S2).

For MW calculations, we used amino acid sequences (Dataset S3). Further details can be found in the Materials and Methods section.

As GltA and AcnB of the TCA cycle are severely restricted and TCA cycle is associated with energy generation, we first determined the ATP producing modules of Ngo. ATP can be produced in three different modules: substrate-level phosphorylation of glucose, acetate overflow metabolism, and TCA cycle. Reactions associated with each module that are carrying flux can be found in Dataset S4. Next, we require the cellular protein content for Ngo. Since this data is also unavailable, we minimized the total protein content, such that the predicted growth rate matches closely to experimental observation, 0.73 *h*^−1^ (D. *et al*. 2023). From the optimization problem, the minimum protein content (Φ) found was 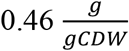. This value is in par with the protein content of *E. coli*, which is 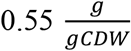 (B., M. and E. 2021) Additionally, we connected ATP synthase

(ATPS) with the TCA cycle with an additional constraint, ATPS can carry 65% of the flux of succinate dehydrogenase (SDH).

With the additional constraint, upon simulation of the model with a Φ of 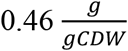, we inspected the activity of each ATP generation modules. ATP generation through substrate-level phosphorylation kept increasing with increasing glucose uptake. Interestingly, ATP generation through acetate overflow does not begin until glucose uptake rate hits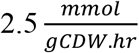, at which growth rate reaches 0.67 *h*^−1^ (Figure 2 A). To support increased growth rates, due to high proteome-usage efficiency, additional ATP production through acetate overflow is a well-understood phenomenon in *E. coli* and yeast (Chen and Nielsen 2019). Next, we represented the active part of the TCA cycle through SDH. The flux through SDH kept increasing until the glucose uptake rate hits 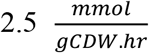 (Figure 2 B). After that, Ngo generated additional ATP through acetate overflow, indicating a reduction of the TCA cycle capacity to produce additional ATP. Surprisingly, right at the reduction of the TCA cycle capacity, the flux of the GltA and AcnB started to decrease drastically compared to other reactions of the TCA cycle (Figure 2 C and 2 D). This indicates, in Ngo, inactivation of the first three steps of the TCA cycle is directly related to the proteomics capacity of the TCA cycle to produce ATP. At the experimentally observed growth rate (0.73 *h*^−1^), first three steps of the TCA cycle were almost completely suppressed, and ATP generation was dominated by the TCA cycle, with a significant contribution from acetate overflow and substrate-level phosphorylation of glucose (Figure 2 E).

**Figure 2.**
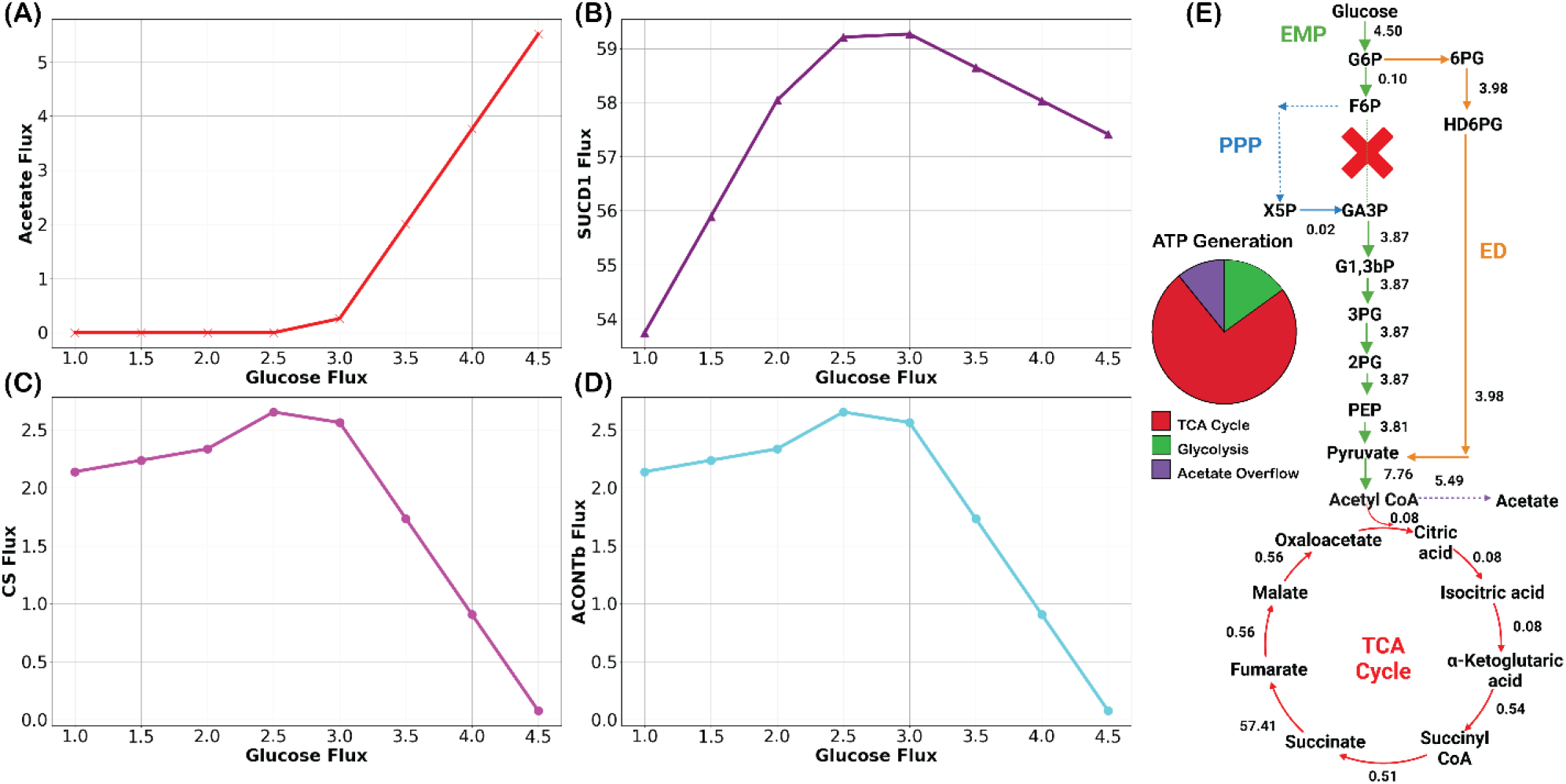
Metabolic flux responses to low protein content 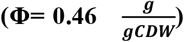 in Ngo. (A) Early onset of acetate secretion indicates enhanced overflow metabolism. (B) Succinate dehydrogenase flux variation. (C) Reduced GltA and (D) AcnB fluxes under protein scarcity. (E) Constrained TCA cycle flux 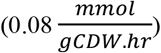 increases ATP via acetate overflow (E). “X” represents the absence of PFK. Additionally, the reaction fluxes are in 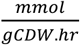.

### Increased Protein Availability Send more Flux through GltA and AcnB

To further test the hypothesis that proteome allocation plays the decisive role in restricting the first three steps of the TCA cycle, we increased the Φ of Ngo to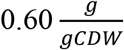. If the hypothesis is accurate, three aspects will be noticed in the Ngo metabolism. First, the onset of acetate secretion will be delayed, as more protein is available for investment in the TCA cycle. Second, the bulk of the ATP will be generated from the TCA cycle and the contribution of substrate-level phosphorylation will remain similar. Third, the first three steps of the TCA cycle will carry much higher fluxes compared to the low Φ.

Indeed, all three aspects were observed with the Φ of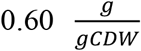. The substrate-level phosphorylation kept increasing with the increased glucose uptake same as observed in low protein content case. The onset of the acetate overflow delayed until 0.82 *h*^−1^ (final growth rate 0.86 *h*^−1^) (Figure 3 A). The diminishing SDH flux matched with the onset of acetate production (Figure 3 B). Finally, GltA and AcnB of the TCA cycle carried much higher fluxes compared to the lower protein content and contribution of the acetate overflow in ATP production was minimum (Figure 3 C, 3 D, and 3 E). To quantify that, we normalized the fluxes of the first three steps of the TCA cycle with final biomass growth rate. For lower Φ, the normalized flux of first three steps of the TCA cycle is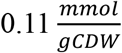, whereas the normalized flux is 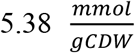 for the higher Φ —a substantial increase by a factor of 47. In contrast, the biomass increase in the higher Φ case was 1.18 times only (0.73 *h*^−1^ in the low and 0.86 *h*^−1^ in the high protein content).

**Figure 3.**
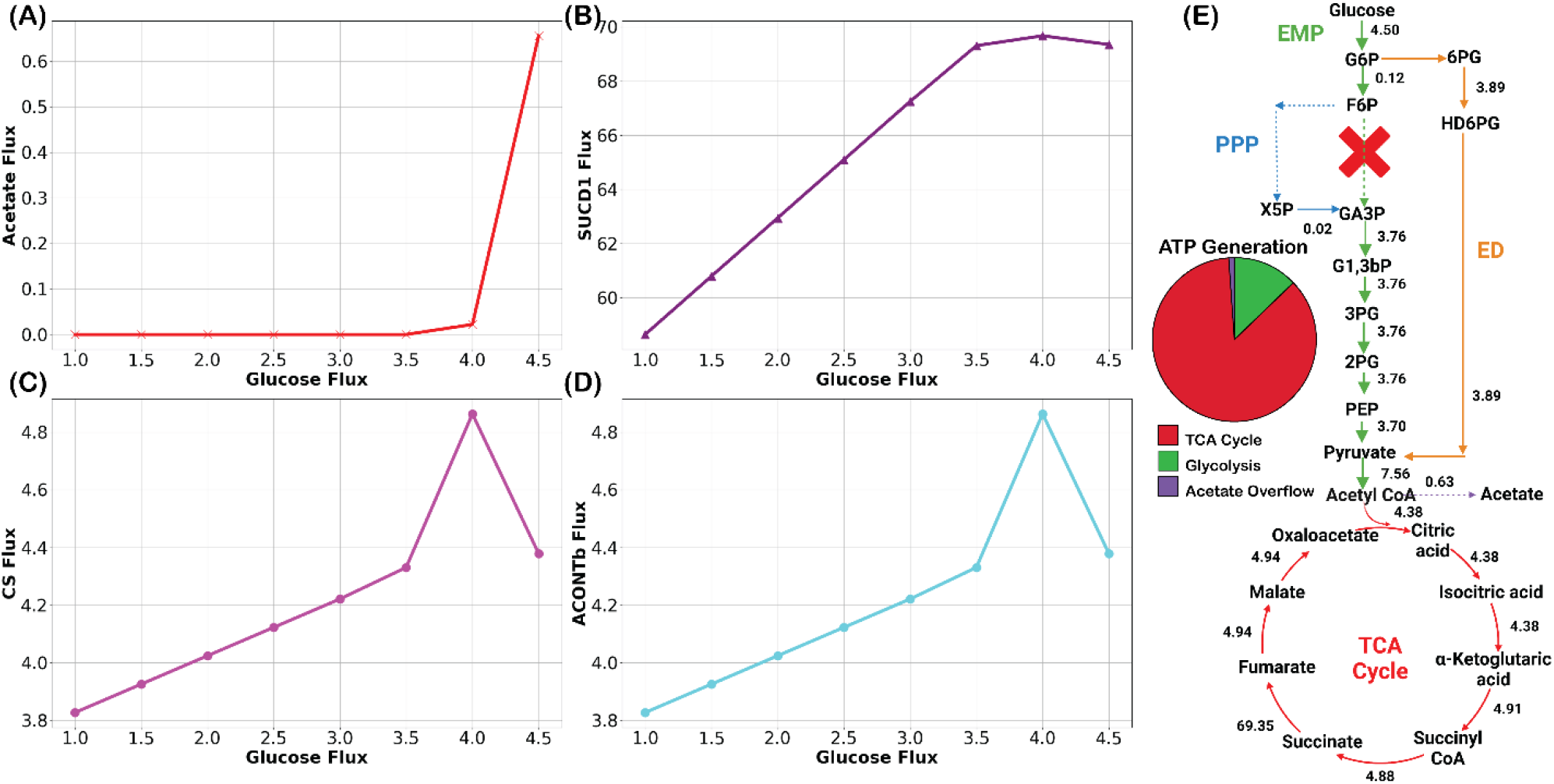
Metabolic flux responses to high protein content 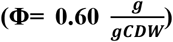. (A) Delay of acetate secretion indicating reduced overflow metabolism. (B) Variations in succinate dehydrogenase flux with glucose uptake. (C) Elevated GltA and (D) AcnB fluxes under high protein availability. (E) Increased flux through the initial TCA cycle steps 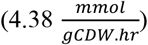 correlates with negligible ATP production via acetate overflow. (E). “X” represents the absence of PFK. Additionally, the reaction fluxes are in 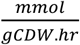.

In conclusion, we conclusively demonstrated that optimal protein allocation strategy can successfully predict experimentally observed metabolic phenotype of Ngo. Our findings will open new possibility to understand the metabolic niche of pathogenic bacteria from the optimal protein allocation perspective and contribute towards global health sustainability.

## Authors’ contributions

Conceptualization, N.S. and N.B.C.; methodology, N.S. and N.B.C.; data analysis, N.S. and N.B.C.; investigation, N.S. and N.B.C.; resources, R.S.; coding, N.S. and N.B.C.; drafting of the manuscript, N.S. and N.B.C.; writing: original draft preparation, N.S. and N.B.C.; writing: review and editing, N.S., N.B.C., and R.S.; supervision, R.S.; funding acquisition, R.S. All the authors have read and agreed to the published version of the manuscript.

### Conflict of interest statement

The authors declare no conflicts of interest.

## Funding

We gratefully acknowledge the funding support from the National Institute of Health (NIH) R35 MIRA grant (5R35GM143009) and National Science Foundation (NSF) CAREER grant (1943310).

## Data Availability

All codes have been deposited in GitHub (https://github.com/ssbio/Gc_protein_constrained).

## Notes

https://github.com/ssbio/Gc_protein_constrained

### Competing Interest Statement

The authors have declared no competing interest.

